# Novel Amplicons based Method for Near-Full-Length Genome (NFLG) Sequencing of HIV-1 Group M Recombinant Forms

**DOI:** 10.1101/2024.08.04.606545

**Authors:** Roy Moscona, Tali Wagner, Miranda Geva, Efrat Bucris, Oran Erster, Neta S. Zuckerman, Orna Mor

## Abstract

Over the years the spread of HIV-1 across the globe resulted in the creation of multiple subtypes and new recombinant forms (RFs). While the pol gene region of the HIV-1 genome is used for resistance mutations analysis and initial detection of RFs, whole genome sequencing analysis is required to determine recombination events across the viral genome. Here, we present a novel robust near-full length genome (NFLG) sequencing approach for the sequencing of HIV-1 genomes, out of clinical whole blood samples. This method has been successfully tested for various HIV-1 subtypes and RFs.

The method is based on an in-house developed set of 32 pan-genotypic primer pairs, divided into two pools, each containing 16 primer pairs covering the entire HIV-1 genome. Two parallel multiplex PCR reactions were used to generate 32 overlapping DNA fragments spanning the HIV-1 genome. Nextera XT protocol was used to obtain barcoded DNA libraries, which were sequenced with the Illumina Miseq platform using a V3 kit. A consensus sequence was determined for each sample and was used to define recombination events across the genome. For this aim, a combined analysis of several computational tools including HIV BLAST, phylogenetic analysis, RIP, SimPlot++ and jpHMM were employed. Overall, plasma samples from 33 patients suspected to carry RFs and 2 different, known pure subtypes controls, were included in this study. Genome coverage varied between RFs, while the gag and pol genes were nearly fully covered, the highly variable env gene region was not. Yet, these NFLG analyses enabled the identification of recombination events genome wide.

In summary, we describe a methodology for HIV-1 NFLG sequencing, which is based on partially overlapping, multiple PCR fragments, spanning the HIV-1 genome. Additionally, this newly refined method was compared to HIV-1 NFLG results of PCR-free metagenomic sequencing and proved to obtain greater coverage of the HXB2 reference genome. Yet, further testing and validation on a larger cohort is required. Still, this method enables sequencing of 20 different patient samples in a single MiSeq sequencing run and was used for the characterization of different HIV-1 RFs and pure subtypes circulating in Israel.

## 1. Introduction

The global HIV-1 pandemic can be attributed to HIV-1 group M (major). Over the years, as the HIV pandemic spread globally within the human population, an increase in viral genetic diversity has been generated, creating a split of group M into several different viral subtypes (A–D, F–H, J, K, L). The leading factors for viral diversity are the high mutation rate of the virus and recombination events, including intra and inter subtypes, which lead to the creation of new recombinant forms (RFs) [1], [2]. Circulating recombinant forms (CRFs), are RFs identified in at least three epidemiologically nonrelated individuals, while unique recombinant forms (URFs), are patient specific.

Since sequencing of the *pol* gene of the HIV-1 genome (mainly the protease {PR}, reverse transcriptase {RT}, and integrase {INT} genes) is still highly recommended before starting anti-retroviral treatment (ART) treatment, many studies utilize these sequences to define RFs [3], [4]. However, regions other than the *pol* gene are not routinely included when analyzing viral sequences, obtained from whole blood samples. For the characterization of recombination events across the viral genome, full or near full-length genome (NFLG) sequencing methodology are utilized. Commonly, PCR amplified DNA fragments, varying in size and length are produced to cover the entire viral genome. These amplicons serve as input for sequencing, either Sanger or Next-Generation Sequencing (NGS). Alternatively, metagenomics, which do not require amplification of the viral genome, could be used. However, the latter demands high resources per sample and is not feasible for large scale clinical settings. Previously, several different methods for HIV-1 NFLG sequencing were developed, some also capture RFs [5]–[8].

Here, we present a novel robust, relatively simple approach, for a pan HIV-1 NFLG sequencing protocol based on multiple short PCR products. This method was developed to capture the clinically relevant HIV-1 group M pure subtypes and CRFs circulating in Israel and in other developed countries. To validate our novel method, we compared our NFLG results to the sequencing results of a PCR-free metagenomic method.

## 2. Materials and Methods

### 2.1 Sample selection

In a previous study by the research team of the National HIV Reference Laboratory (NHRL), HIV-1 CRFs and URFs in Israel were defined, based on HIV-1 PR and RT partial sequences for people living with HIV-1 (PLWH) diagnosed between 2010 and 2018 [3]. Here, samples from 33 patients suspected of representing different CRFs and URFs (and another 2 samples of known HIV-1 pure subtypes served as control, totaling in 35 samples), were used for the assessment of a new NFLG sequencing method.

### 2.2 Selection of primers for multiplex PCR

The LANAL database (https://www.hiv.lanl.gov/components/sequence/HIV/search/search.comp) was searched for full HIV-1 sequences (sequence length >= 9000 bp) that were sampled between 2000-2019 aiming to generate a diverse input dataset for primer selection. To narrow down the search results, only HIV-1 subtypes and RFs known to be circulating in Israel were selected. These included different HIV-1 group M pure subtypes (B, C, A1, D, G, A6, F1, H, U) and RFs (AE, AG, CRF_206, BC, A6B, A5U) and complex forms (cpx) (06, 45, 27). The query resulted in 291 HIV-1 genome sequences, of which 118 representative sequences for each HIV-1 pure subtype, RF, cpx (above-mentioned) were selected. This dataset was further divided into two sets of sequences: a training set and a testing set. The training set contained newer isolates, collected between the years 2003-2019 (n=76), while the testing data set contained older sequences isolated between the years 2000-2002 (n=42). The training dataset served as a template for designing new primers, while the testing dataset was utilized virtually, for *in silico* PCR analysis, to ensure the designed primers were suitable, before synthesizing the primers for *in vitro* multiplex-PCR.

The generated dataset of HIV-1 sequences was aligned to the HXB2 reference genome by Geneious Prime (Dotmatics, Boston, MA, version 2022.2.2, Build 2022-08-18) default mapper using default settings. The primal-scheme [9] tool (https://primalscheme.com/) was used to identify conserved regions across the HIV-1 genome in a sliding windows of ∼400bp (Fig. 1). Primers were constructed to enable amplification of the whole viral genome in two separate PCR reactions divided into two pools of overlapping fragments, varying in length with an average fragment size of 368bp. The primal-scheme primers design was further edited using Geneious Prime, to incorporate ambiguous nucleotides and adjust the genomic location of each primer, whilst keeping optimal biochemical features and maintaining an appropriate distance between PCR product fragments yet keeping some overlap between them. The two pools of primers and their locations in HXB2 are presented in Supp. Table 1.

**Fig. 1.**
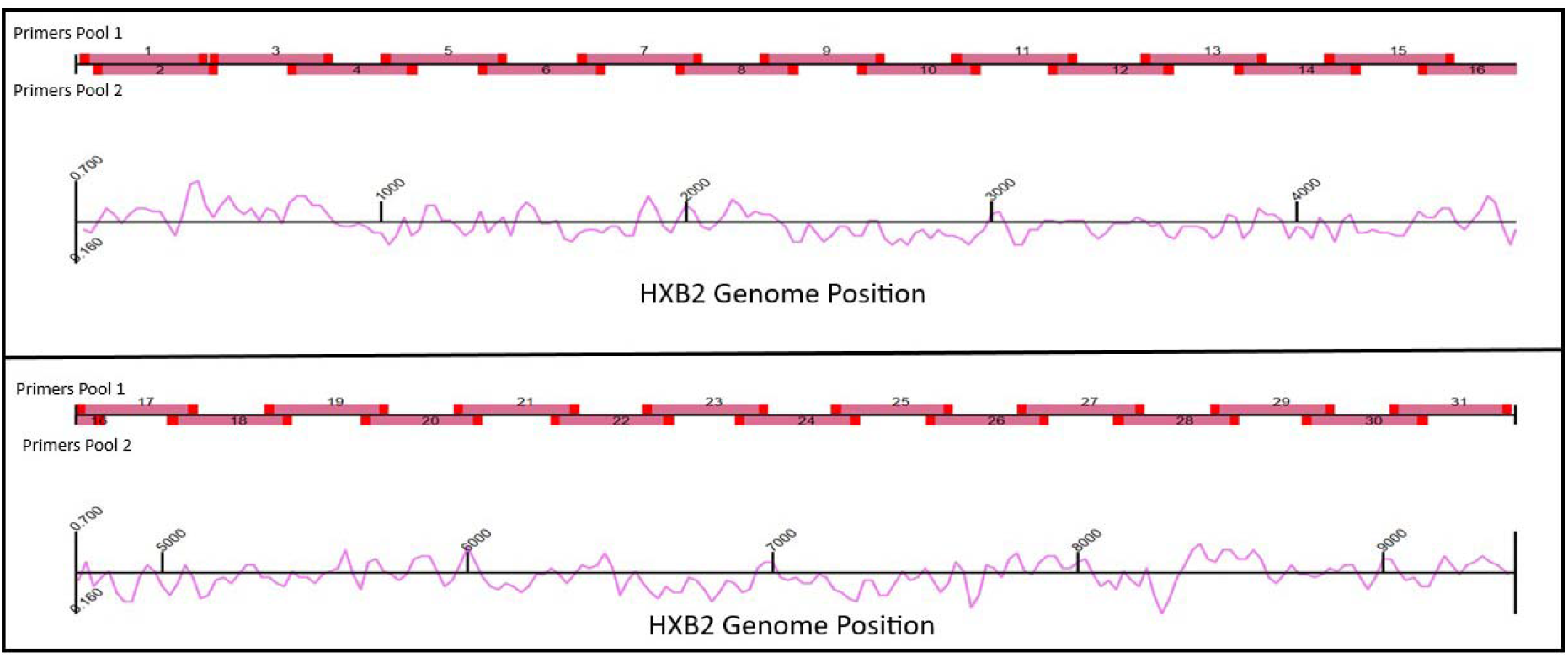
Schematic mapping of the primers covering HIV-1 HXB2 genome. The general outline of the primers across the HIV-1 reference HXB2 genome designed using primal-scheme [9] is shown. The primer positions are colored in bright red; the amplicons are shown in darker shaded red.

### 2.3 HIV-1 amplification and DNA Library preparation for NGS

For the novel amplicons-based HIV-1 NFLG, first HIV-1 RNA was extracted from plasma of whole blood samples using Nuclisense protocol (BioMerieux), complementary DNA (cDNA) was synthesized using SuperScript III first strand kit (Thermo Fisher Scientific, Waltham, MA). Two multiplex-PCR reactions were performed in parallel in two test tubes, each containing a different primer pool (pool 1 or pool 2). Next, the two pools were combined in equal molarity and diluted as required according to the NexteraXT protocol (Illumina, San Diego, CA).

For the metagenomics (virome sequencing) protocol four samples were selected and HIV-1 RNA was extracted as described before. However, the key difference here was the cDNA synthesis, SuperScript III first strand kit (Thermo Fisher Scientific, Waltham, MA) was used. Followed by, SMARTer Stranded RNA-Seq Kit (Takara Bio, San Jose, CA), that was used to generate un-amplified cDNA library.

For both methods (amplicons and metagenomics), the resultant libraries were purified with AMPureXP beads (Beckman Coulter, Brea, CA). Library concentration was measured by Qubit dsDNA HS Assay Kit (Thermo Fisher Scientific, Waltham, MA). Library validation and mean fragment size (∼300bp) was measured by TapeStation4200 (Agilent, Santa Clara, CA) with DNA High-Sensitivity D1000 (Agilent, Santa Clara, CA). Following dilution to 4nM equal molar quantities of the libraries were pooled, denatured and diluted to final concentration of 15pM. The final DNA library was subjected to paired-end sequencing (2×100bp) either on NovaSeq (Illumina, San Diego, CA), for metagenomics, or MiSeq (Illumina, San Diego, CA), for the new amplicons-based methodology.

### 2.5 Bioinformatics analysis

FastQ sequences were subjected to a custom pipeline, tailored to analyze viral genome NGS data. This pipeline was used to generate a consensus sequence for each sample and it has been described in (https://github.com/NetaZuckerman/UPv/blob/main/pipelines/generalPipeline.py).

### 2.6 Analyses of HIV-1 recombination forms with phylogeny, HIV-BLAST, RIP, SimPlot++ and jpHMM

Phylogenetic trees were constructed with the neighbor-joining algorithm, while implementing a 1000-bootstrap repetition, used for branch assessments. Four different phylogenetic analyses were performed. First, a phylogenetic tree based on NFLG sequences was built. This phylogeny approach enabled the characterization of the dominant viral clade of recombination (major parent). Next, an additional three independent phylogenetic trees were constructed, one for each viral gene: *gag, pol, env*. This approach was chosen to identify specific recombination events in the main three viral genes, aiming to identify minor parents. All phylogenetic trees were constructed with known pure subtypes and RFs as reference sequences. This enabled the classification of each sample to an appropriate HIV-1 viral clade. HIV-BLAST was used to assess the similarity to known public repository sequences. Furthermore, Simplot++ was used to identify genome wide recombination events [10]. Simplot++ is utilizing similarity plots to determine the probability of the recombination event and its breaking points across the HIV-1 genome. Simplot++ was used in BootScan mode with a 100-bootstrap repetition, using a 400 bp window with 20 bp step size with the TN93 model for substitutions.

Additionally, jumping profile hidden Markov model (jpHMM) [11] and recombinant identification program (RIP available at www.hiv.lanl.gov/content/sequence/RIP/RIP.html) methods were used with default parameters. Final characterization of each CRF or URF was based on a combinatory analysis of the results obtained by all analytic tools used here, while considering each tool’s advantages and drawbacks in resolving recombination events across the viral genome.

## 3. Results

### 3.1 Comparison between metagenomics and the novel amplicons method for HIV-1 NFLG sequencing

To compare the sequencing results obtained by the newly developed amplicons method to un-amplified metagenomics (virome sequencing), four samples were selected. While the average total number of single-end reads per sample obtained by the metagenomics approach was 66,178,474 (range: 57,842,929-71,825,644) reads, only an average of 59,560 reads (0.09% of total reads) were mapped to the HIV-1 HXB2 reference genome. On the contrary, for the novel amplicons NFLG method, only 2,588,962 (range: 1,254,560-3,099,939) total reads were obtained per sample, however on average 2,191,779 reads (84.6%, of total reads) were mapped to the HIV-1 HXB2 reference genome. Comparison of the coverage (>5 reads per position in both methods) across HIV-1 HXB2 reference genome between the metagenomics and the NFLG amplicons methodology, revealed that the amplicons displayed a higher total coverage when directly compared. The two sequencing methods produced identical results, while surprisingly the amplicons managed to reach greater coverage and depth, sequencing new positions across the HXB2 genome, which were not reported by metagenomics (Fig. 2). Generally, among the different HIV-1 subtypes, metagenomics obtained an average coverage of 85.4% while the newly developed amplicons approach covered 90.6% of the HXB2 reference genome bases. The difference in coverage observed between samples could be attributed to diversity in viral clades. The coverage of the control sample of HIV-1 pure subtype B was 97.27% vs 95.38%, amplicons vs metagenomics, respectively, (sample 99666, Fig. 2A), while the coverage of the other control sample of HIV-1 pure subtype C (sample 100545, Fig. 2B) was lower (88.73% vs 81.94%). For the other two non-pure HIV-1 subtype samples, the same range was observed; sample 84808 (Fig. 2C, 89.3% vs 74.6%) and sample 79455 (87.65% vs 73.77% Fig. 2D), amplicons vs metagenomics, respectively. When aligning the HIV-1 NFLG consensus sequences generated by the two methods (amplicons and metagenomics), a full match was observed in common positions in all samples.

**Fig. 2.**
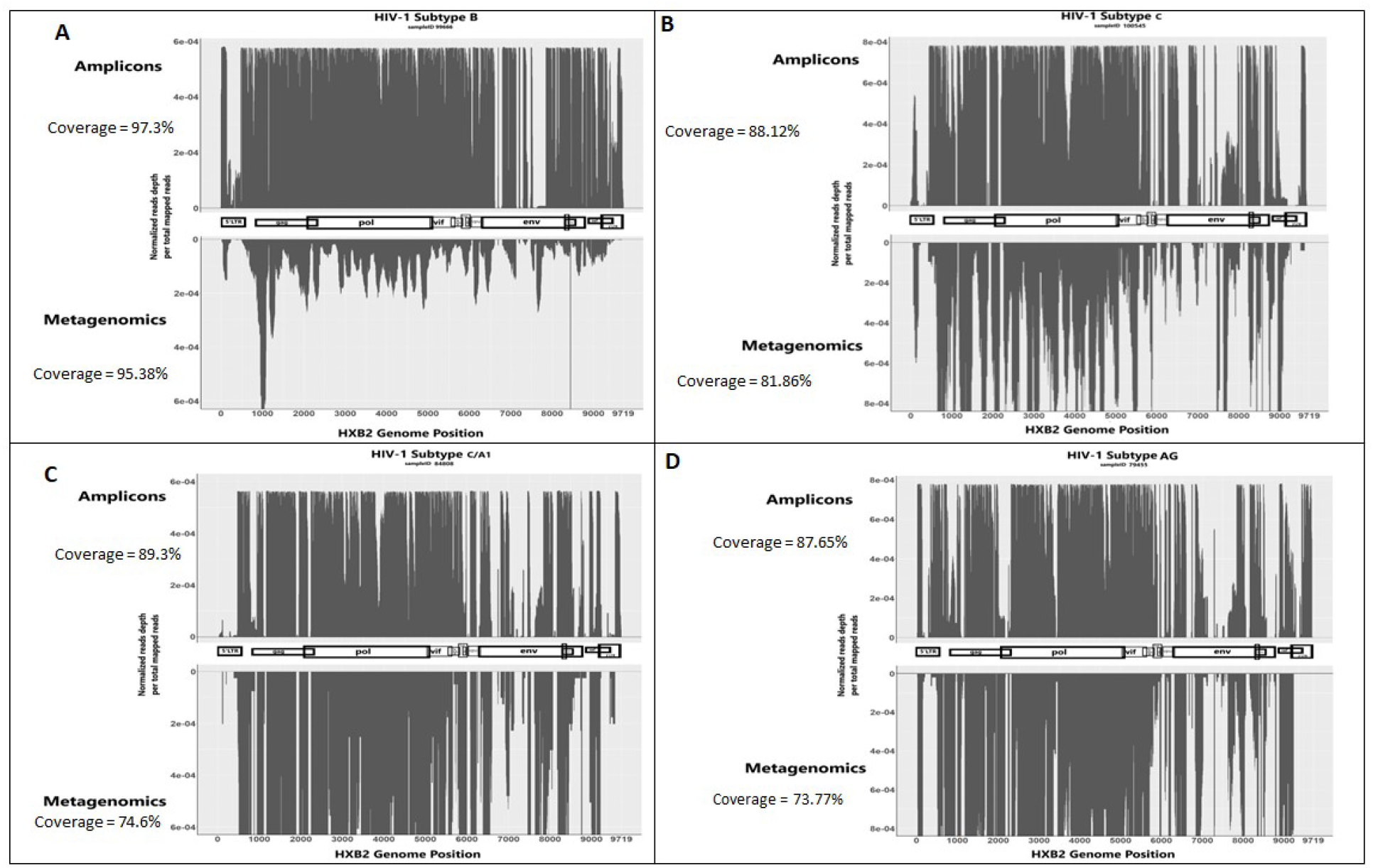
Comparison of HIV-1 HXB2 genome coverage between the novel amplicons method and metagenomics. Values (Y-axis) represent normalized number of reads mapped to the corresponding base position in the HXB2 genome (X-axis). Y-axis directionality (+) for Amplicons and (-) for Metagenomics has no mathematical meaning and was plotted for comparison of each genomic position between the two methods. In the middle of each plot the HXB2 genome map is depicted with genomic regions/genes as follows: 5’LTR, gag, pol, vif, vpr, vpu, env, nef, 3’ LTR.

### 3.2 In silico down sampling scaling to resolve the minimal number of reads required for maximal coverage by the novel amplicons method

The coverage of bases of the HIV-1 reference genome HXB2 varied between different HIV-1 subtypes. Thus, to enable a unify criteria for a pan-HIV-1 settings, a calculation was made to determine the optimal number of reads required for maximal coverage of the 9,719 bases of the HXB2 reference genome, while sequencing the maximal number of samples in parallel on a single run of a Miseq V3 kit. For the down-sampling analysis only 10 samples were sequenced in parallel, and each sample received an even number of reads. We concluded that in the tradeoff between the number of reads assigned for each sample and coverage of HXB2 bases, the required number of paired-end reads is between 5 × 10^5^ to 7.5 × 10^5^ reads, as there was little increase in coverage beyond 750K reads (Fig. 3). Beyond 7.5 × 10^5^ reads, additional reads did not increase the coverage significantly. Hence, we concluded that one Miseq V3 kit was sufficient to run 20 samples in parallel, while keeping adequate coverage and maximal usage of samples sequenced per kit.

**Fig. 2.**
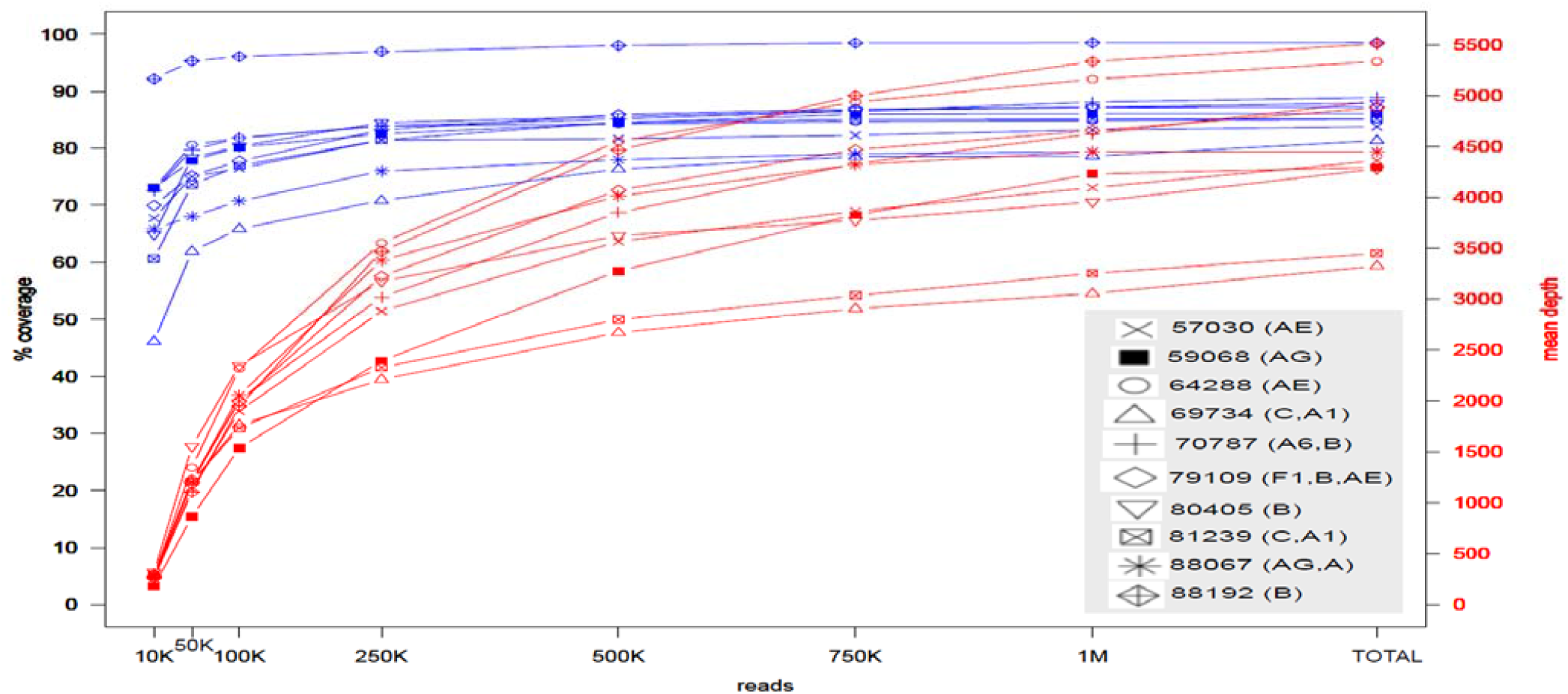
Reads down-sampling analysis for establishing the optimal number of reads needed to allocate for each sample. The blue lines correspond to the Y-axis on the left, representing HXB2 genome precent coverage. The red lines correspond to the Y-axis on the right corresponding to the mean sequence depth (in number of reads). The X-axis are the number of reads (K=thousand, M=million) randomly sampled. For each one of the 10 samples analyzed, a shape was assigned as shown on the grey legend, also in the legend in parenthesis next to sample number is the assigned HIV-1 subtype. The top blue line with diamond shape corresponds to sample 88192 of HIV-1 subtype B, while bottom blue line with triangle shape corresponds to sample 69734 of HIV-1 subtype C and A1. In the plot X-axis, the TOTAL average number of mapped paired-end reads was 1,095,890 and varied by samples and subtypes.

### 3.3 Identification of RFs based in the NFLG sequences

To resolve the recombination events across HIV-1 NFLG sequences for each sample, several analytic tools were practiced. The tools applied here to identify RFs were phylogenic analyses, HIV-BLAST SIMPLOT++, LANL RIP, jpHMM [3], [8], [10], [11]. The subtyping tools displayed a general strong agreement between themselves, identifying a similar if not identical HIV-1-subtype for every sample. To increase the confidence level that each recombination event reported by each tool is not an artifact, we sought agreement between at least two different tools, to increase the confidence level in the resolved recombination event.

### 3.4 Resolution of recombination events across HIV-1 NFLG sequences

For each sample a combinatory analysis was used to reach a final conclusion that would determine which of the HIV-1 subtypes underwent recombination (Table 1). Samples 57030 and 64288 were classified in all four tools as RF AE and resolved to be CRF01_AE. Both samples had subtype B classification in only one tool: sample 57030 in jpHMM while sample 64288 in the phylogenetic tree, since these observations were not consistent with at least two different tools, we concluded those as artifacts and resolved these samples as RF AE. Sample 79109 had displayed a minor sequence of RF AE, in addition to subtypes B and F1 recombination. This finding was reported by two different tools (jpHHM and Simplot++), hence it was concluded as true positive and this sample was assigned a subtype F1 and B and CRF01_AE recombination. Samples 65699 was resolved as subtype B and C as at least two tools reported the presence of these subtypes. Similarly, sample 89194 had subtypes B and C finding in addition to subtype D reported by three tools including the phylogenetic analyses of the *gag* and *pol* genes, thus it was resolved as subtypes B and CD or 41_CD based on NFLG phylogeny.

**Table 1.**
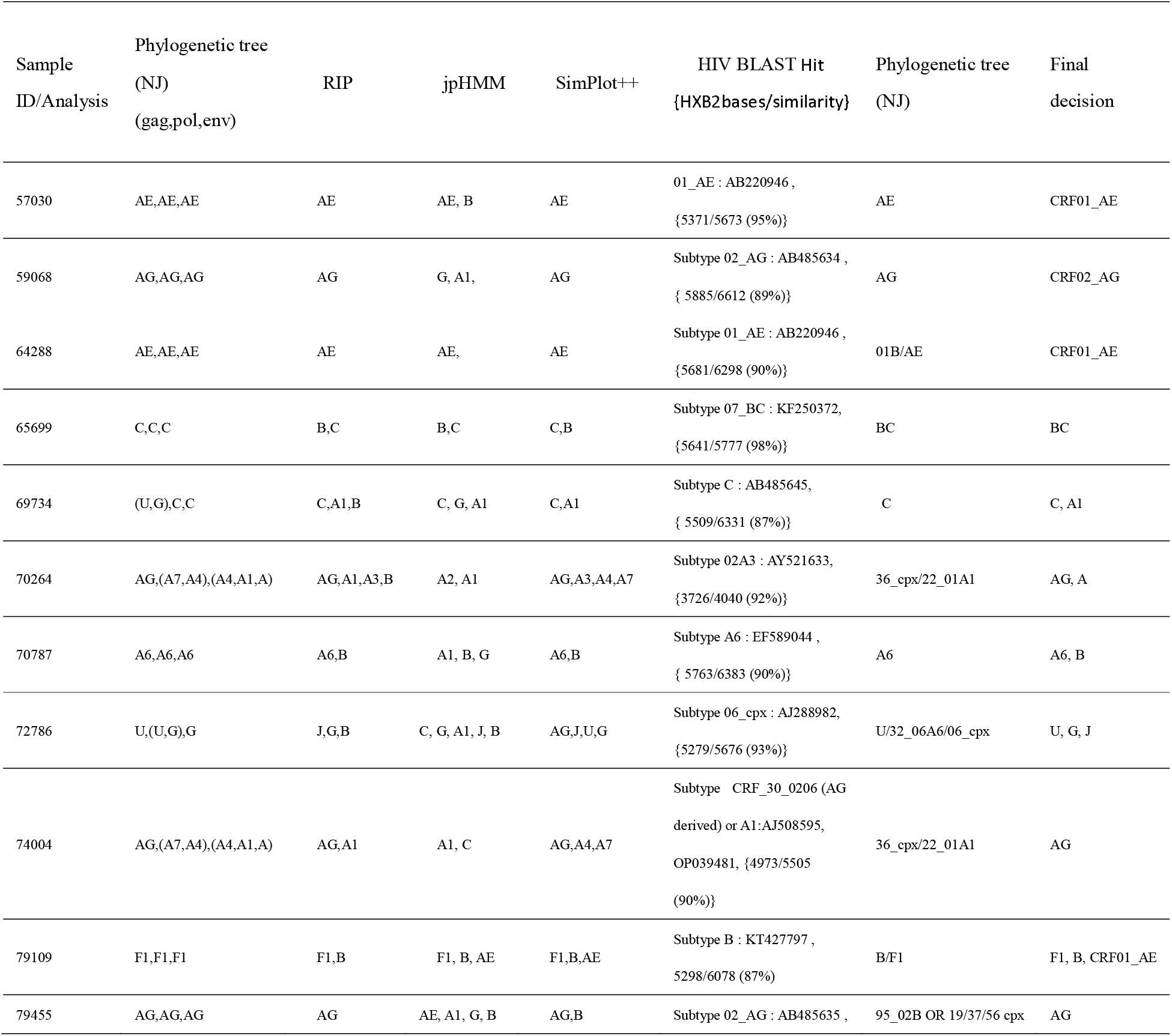
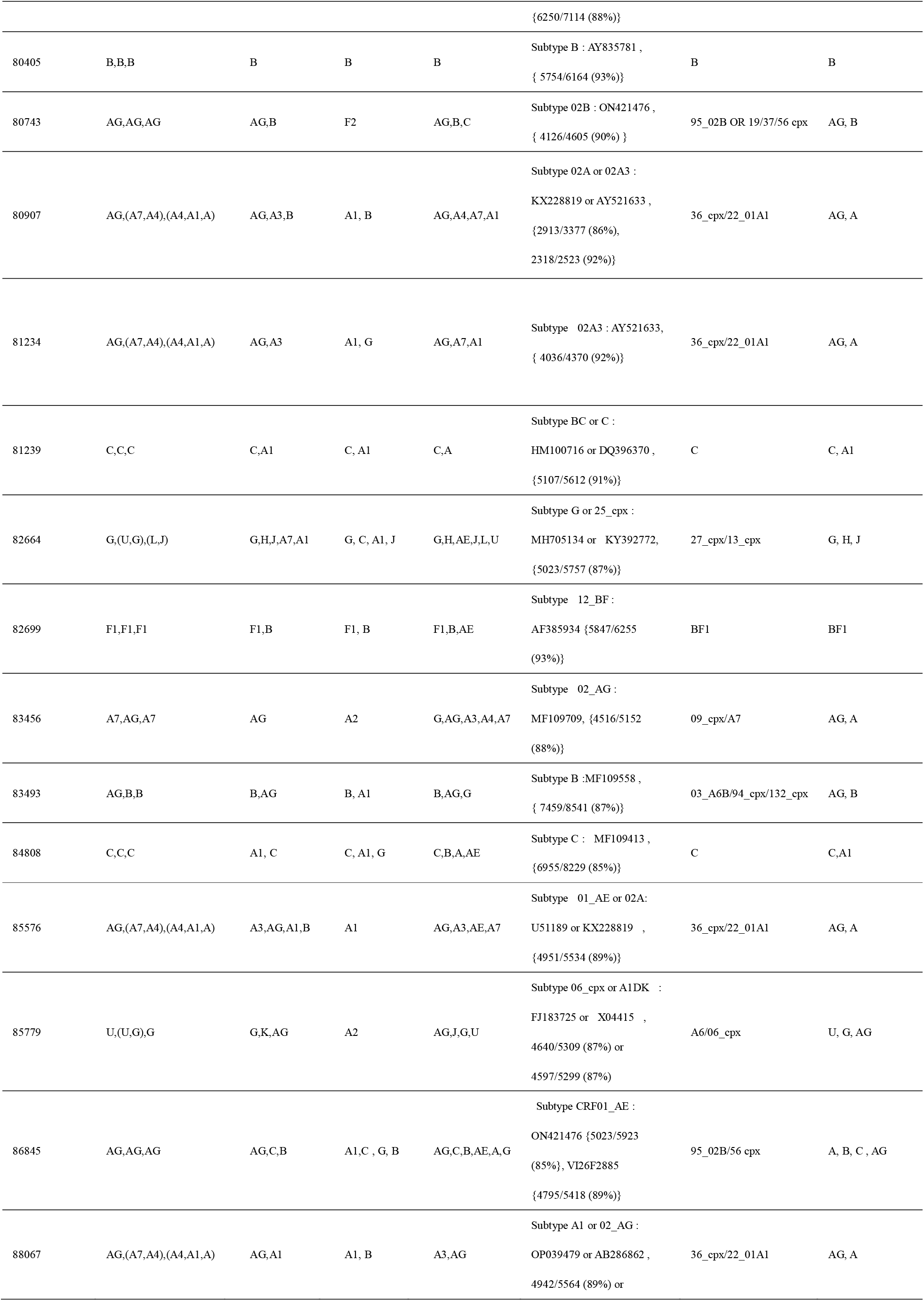
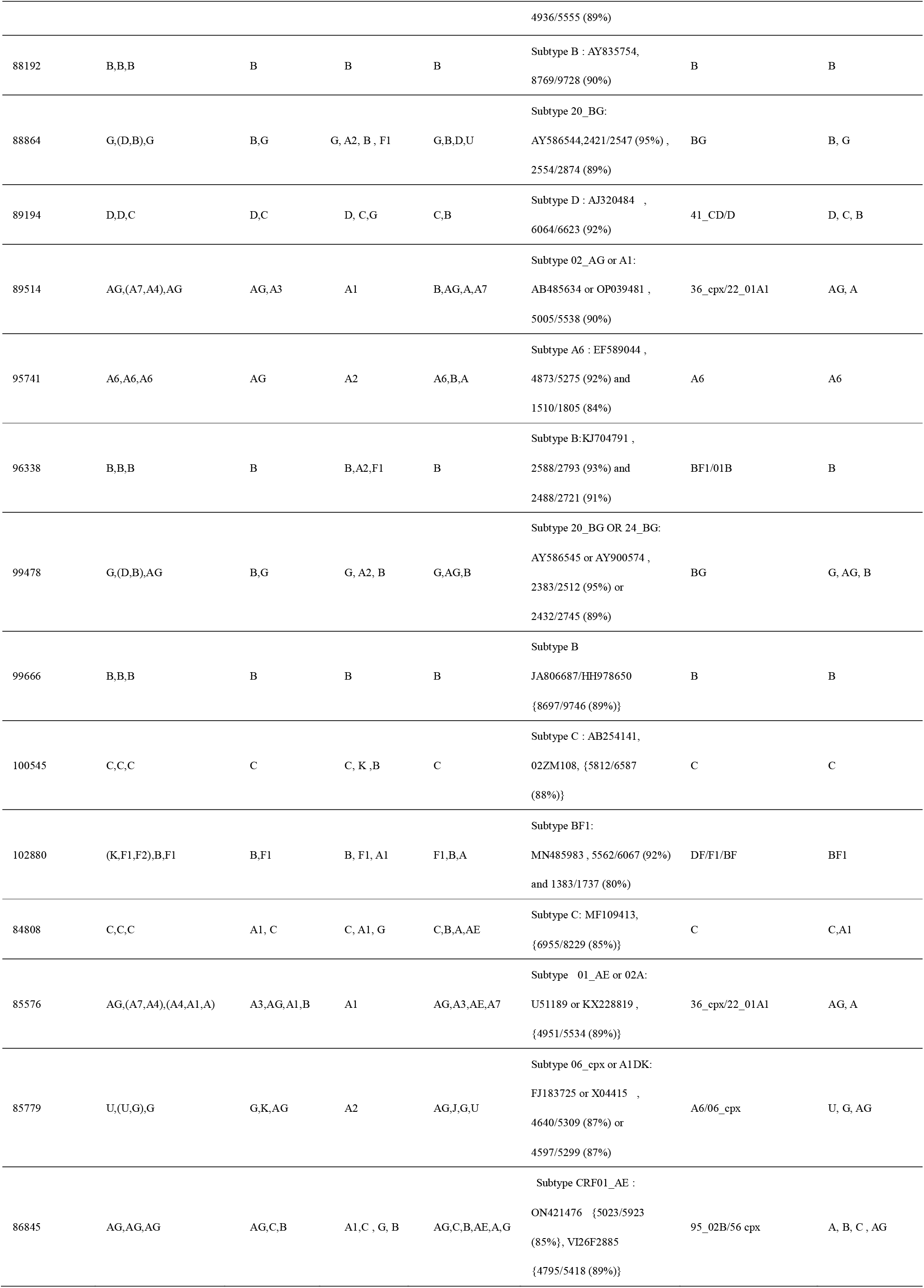
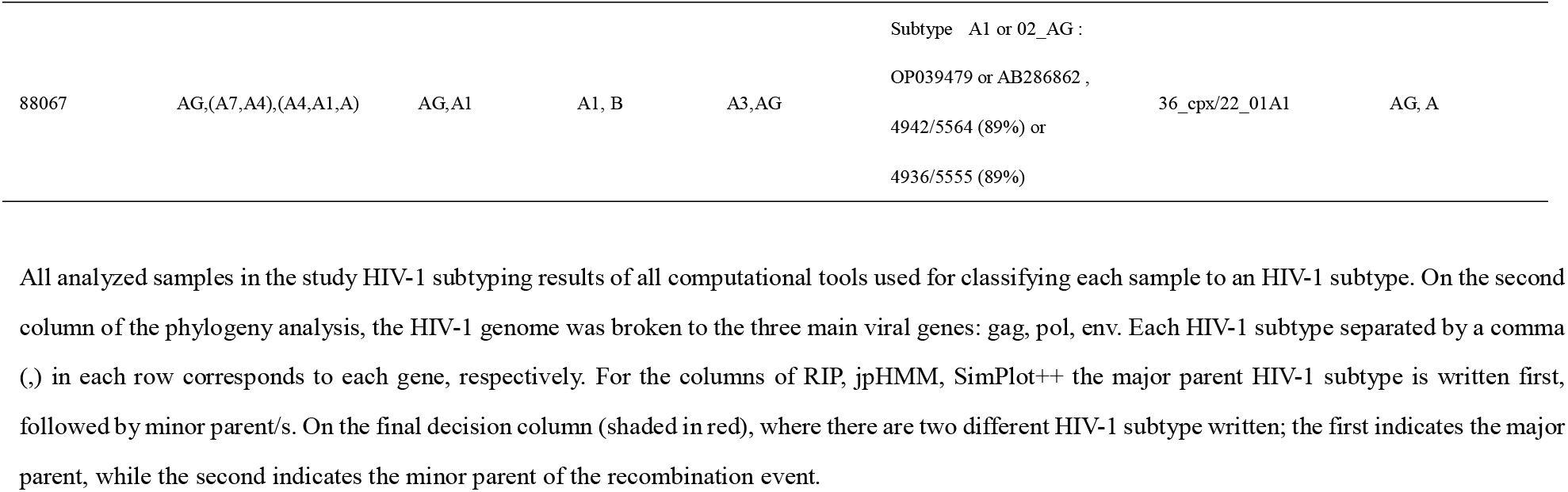
Combinatory analyses for the resolution of HIV-1 subtypes recombination across the study samples.

#### 3.4.1 Table

Samples 69734, 81239, 84808 were resolved as recombination of subtype C and A1, as at least two tools displayed that identification, however the NFLG phylogeny and HIV-BLAST favored subtype C, since it was the major parent in those samples, thus we concluded a subtypes C and A1 recombination. Samples 59068, 74004, 79455 were resolved as either RF AG or CRF02_AG, as all methods (except jpHMM) agreed with that classification. Samples 70264, 80907, 81234, 83456, 85576, 88067, 89514 were resolved as RF AG and subtype A, however all (besides one) were suspected to be cpx 36 or 22_01A1, based on NFLG phylogeny. Only sample 83456 showed a different NFLG phylogeny result with resemblance to cpx09 or subtype A7. The HIV-BLAST results for these samples did not cover enough HXB2 bases (most only covered ∼2,000bp) and the classification to subtype A in all of them seemed to match the minor parent, yet sample 88067 had a ∼5,000bp matching for an HIV-BLAST search with subtype A1 and RF AG. Since in most tools RF AG was the dominant RF, it was concluded as the major parent, while subtype A was the minor parent.

The samples that displayed additional recombination to RF AG were 85779, 86845 with subtypes U and G and RF AG or subtypes A, B and C and RF AG recombination, whether these forms are 06_cpx or CRF01_AE (respectively), based on HIV-BLAST results or URF warrens further investigation. Sample 96338 displayed a smaller coverage across the HXB2 genome compared to the rest of the samples, thus we could not conclude if this is a true recombinant form of subtypes B and F1, so it was resolved as subtype B. On the other hand, samples 82699 and 102880 were resolved as subtypes B and F1 recombinants. Interestingly, the *gag* gene of sample 102880 was classified as an out group for a cluster containing subtypes K, F2, F1, displaying the advantages of a gene-by-gene phylogeny analysis and indicating the recombination event happened in the *gag* gene’s location. Sample 70787 was resolved as subtype B and A6 recombination, even though the phylogeny clustered only with subtype A6, RIP and SimPlot++ indicated a presence of subtype B. Therefore, the major parent was resolved as subtype A6, while the minor parent was subtype B. Sample 95741 was resolved as subtype A6, based on phylogeny and HIV BLAST results as the rest of the tools showed discrepancies and were not conclusive. Samples 80743 and 83493 were resolved as a recombination of subtypes B and RF AG, three tools (phylogeny, RIP, SimPlott++) identified parts of subtype B and RF AG in the sequence, although these samples may be a form of cpx (19/37/56) as they clustered among them on NFLG phylogenetic tree. Additionally, sample 88864 was classified as a recombination of subtypes B and G, while sample 99478 showed subtypes B and G and RF AG, recombination. Sample 85576 displayed a complex pattern of different subtypes A recombination among all tools and was suspected to be either 36_cpx or a URF. Similarly, samples 72786 and 82664 displayed a complex pattern of subtypes U, G, J, B, G and RF AG (for sample 72786) and subtypes G, L, U, A1 (for sample 82664), however they may be a form of cpx (06 or 11/13) based on NFLG phylogeny. Samples 80405 and 88192 were resolved as pure subtype B, with no indication of recombination events by any of the tools. Samples 99666 and 100545 served as positive control for pure subtypes B and C, respectively and were resolved as such.

## 4. Discussion

Here we have shown the successful implementation of a novel method for HIV-1 NFLG sequencing. This method was tested with clinical samples of whole blood plasma and shown to be relatively easy to perform in a clinical laboratory setting, while simple and less laborious than previously reported methods for HIV-1 NFLG.

Previous studies have utilized HIV-1 NFLG protocols successfully, some of the previously developed methods include Gall et al. 2012 [6], which developed a pan HIV-1 primer set, which generates 4 overlapping PCR fragments to cover the viral genome, followed by Roche/454 (Life Sciences, Branford, CT) sequencing. This method showed high sensitivity and reproducibility in detecting drug resistance and minor variants, and was able to identify CRF01_AE and CRF14_BG. Moreover, Grossman et al. 2015 [12], [13] have developed a two PCR fragments method that covers the entire HIV-1 genome, however this was tested only on subtypes B and C and RFs CRF01_AE and CRF02_AG. Others, such as Hebberecht et al. 2019 [14] developed a multi-step PCR (RT & Nested) based protocol, generating two amplicons followed by sanger sequencing. Their method managed to identify successfully CRF01_AE and CRF02_AG. Additionally, Lunar et al. 2020 [7] have utilized existing methods for HIV-1 NFLG sequencing to identified 6 new URFs and 3 more potential CRFs, in a country with limited HIV diversity.

Our newly developed method aimed to simplify HIV-1 NFLG sequencing procedure, while capturing pan-HIV-1 genotypic variants, that are circling in developed countries. The approach described here has enabled the identification of HIV-1 RFs including CFRs and potential URFs. However, the method introduced here is lacking full coverage of highly variable regions of the HIV-1 genome like *env* and repetitive regions like the LTR. The amplicons developed method introduced in this pilot study, in optimal conditions, allocating maximal number of reads per sample, surpassed metagenomics in the percentage of HXB2 bases covered, it achieved higher coverage in all 4 compared samples of different HIV-1 pure subtypes and RFs (Fig. 2).

With regards to the *env* gene classification in all used computational tools, the confidence level of that region classification is low, due to lack of full coverage. Consequentially, its classification was difficult to analyze e.g. if a sample had a different subtype classification present only for a relative short number of bases in the *env* gene it was ignored, especially if it did not fit with the rest of the genome classification. Additionally, some analytic tools like jpHMM had a bias towards subtype B assignment for the *env* region. With further regards to jpHMM, this tool does not have RF AG as reference, therefore it classified RF AG as a combination of subtype A (mostly A1) and subtype G, consequently, its classification was ignored with regards to RF AG samples only. On the other hand, HIV-BLAST results should be compared with other methods, as the parts of the HIV genome covered by HIV-BLAST results may have a bias towards the major parent in a RF and parts of the unmatched genome may contain different subtype classification, that could potentially contain the minor parent of the recombination.

## 5. Conclusions

Here, we have successfully simplified the procedure for HIV-1 NFLG sequencing, still future maintenance of the primers designed here will be needed, as new HIV-1 NFLG CRFs and URFs sequences will accumulate over the years in publicly available databases. The primers presented here (Supp. table 1) should be tested on newly sequenced HIV-1 genomes and updated accordingly, to insure maximal capture of ongoing HIV-1 viral diversity with this approach. Overall, our newly developed amplicons approach has proven itself as reliable on this pilot study, yet it should be compared on a sample-by-sample basis with other HIV-1 NFLG methods to test its robustness and on a larger scale cohort and by other laboratories to be fully corroborated as a suitable approach for HIV-1 NFLG sequencing.

## Abbreviations

ART: anti-retroviral treatment
CRFs: circling recombinant forms
INT: integrase
NGS: next generation sequencing
NFLG: robust near-full length genome
NHRL: National HIV Reference Laboratory
PLWH: people living with HIV-1
PR: protease
RF: recombinant form
RT: reverse transcriptase
URF: unique recombinant form

## Supplementary Material

**Supplement table 1.**
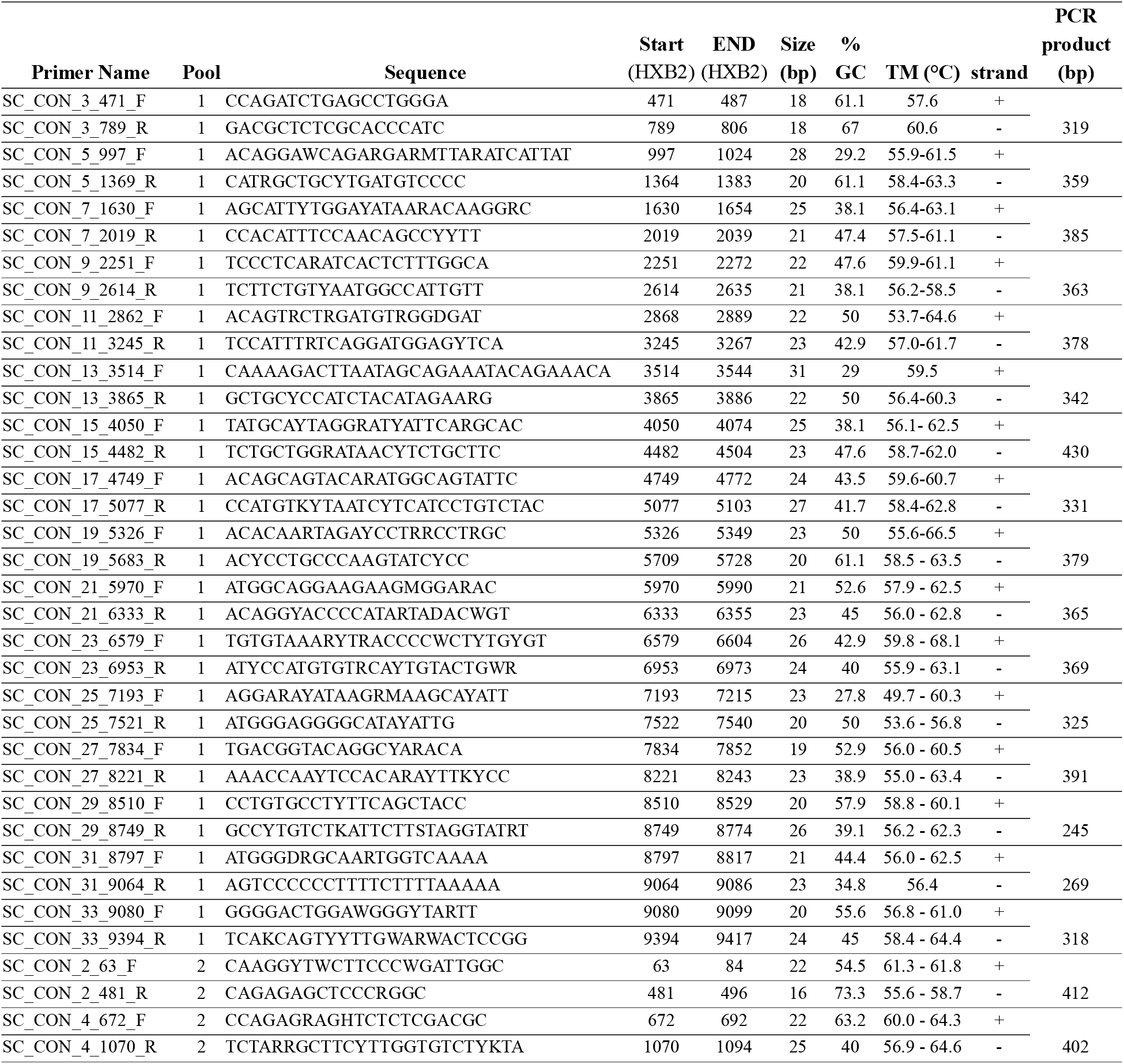
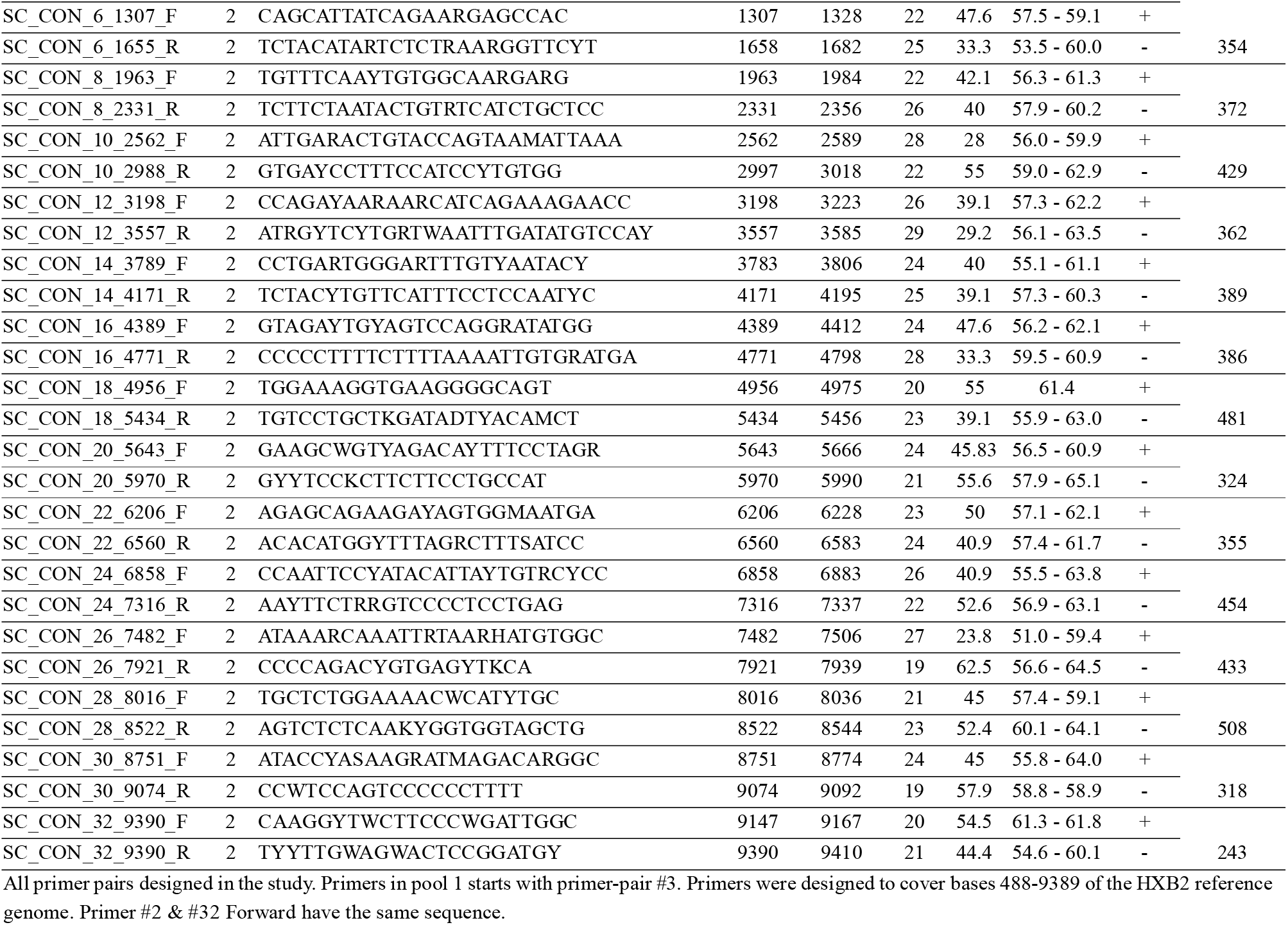
Primers designed in this study to generate the amplicons for HIV-1 NFLG sequencing.

## Availability of Data and Materials

Raw data can be provided upon request from the authors.

## Author Contributions

Conceptualization, O.M. and R.M.; methodology, O.M., R.M., T.W., M.G., E.B., O.E., N.Z.; formal analysis, O.M., R.M., T.W., N.Z.,; resources, O.M., E.B., O.E., N.Z.; writing—original draft preparation, R.M. and O.M.,; writing—review and editing, O.M., R.M., T.W., M.G., E.B., O.E., N.Z.; All authors have read and agreed to the published version of the manuscript.

## Ethics Approval and Consent to Participate

The study was conducted according to the guidelines of the Declaration of Helsinki and was approved by the Institutional Review Board of the Sheba Medical Center.

## Acknowledgment

We would like to acknowledge Zehava Yossefi from the NHRL for her excellent and persistent contribution to maintenance of the NHRL HIV database.

